# Niche separation increases with genetic distance among bloom-forming cyanobacteria

**DOI:** 10.1101/226860

**Authors:** Nicolas Tromas, Zofia E. Taranu, Bryan D Martin, Amy Willis, Nathalie Fortin, Charles W. Greer, B. Jesse Shapiro

## Abstract

Bacterial communities are composed of distinct groups of potentially interacting lineages, each thought to occupy a distinct ecological niche. It remains unclear, however, how quickly niche preference evolves and whether more closely related lineages are more likely to share ecological niches. We addressed these questions by following the dynamics of two bloom-forming cyanobacterial genera over an 8-year time-course in Lake Champlain, Canada, using 16S amplicon sequencing and measurements of several environmental parameters. The two genera, *Microcystis (M)* and *Dolichospermum (D)*, are frequently observed simultaneously during bloom events and thus have partially overlapping niches. However, the extent of their niche overlap is debated, and it is also unclear to what extent niche partitioning occurs among strains within each genus. To identify strains within each genus, we applied minimum entropy decomposition (MED) to 16S rRNA gene sequences. We confirmed that at a genus level, *M* and *D* have different preferences for nitrogen and phosphorus concentrations. Within each genus, we also identified strains differentially associated with temperature, precipitation, and concentrations of nutrients and toxins. In general, niche similarity between strains (as measured by co-occurrence over time) declined with genetic distance. This pattern is consistent with habitat filtering – in which closely-related taxa are ecologically similar, and therefore tend to co-occur under similar environmental conditions. In contrast with this general pattern, similarity in certain niche dimensions (notably particulate nitrogen and phosphorus) did not decline linearly with genetic distance, and instead showed a complex polynomial relationship. This observation suggests the importance of processes other than habitat filtering – such as competition between closely-related taxa, or convergent trait evolution in distantly-related taxa – in shaping particular traits in microbial communities.

## INTRODUCTION

Microbial communities are composed of potentially diverse groups of lineages, which must be sufficiently similar to survive in the same habitat, yet sufficiently dissimilar to occupy distinct ecological niches and avoid competition. This tension between selection for common traits to survive in a common environment (habitat filtering) and selection for divergent traits to reduce competition (niche partitioning among closely-related species) was recognized by Darwin, and the relative impacts of the two processes on communities are still debated (Cavender-Bares *et al.*, 2009). A pioneering study of microbial communities using phylogenetic marker genes found evidence for phylogenetic clustering, suggesting the importance of habitat filtering in selecting for closely-related taxa sharing specific traits allowing them to survive in a given habitat (Horner-Devine and Bohannan, 2006). However, phylogenetic overdispersion (the opposite pattern as phylogenetic clustering) has also been observed, suggesting that competition between closely-related taxa can lead to niche partitioning (Koeppel and Wu, 2014). Importantly, the power to detect phylogenetic overdispersion depends on the phylogenetic resolution (*e.g.* whether operational taxonomic units are defined at 97, 98 or 99% identity in a marker gene) (Koeppel and Wu, 2014).

Beyond searching for phylogenetic patterns of clustering or overdispersion, explicitly considering the associations between microbial traits and niches can help understand the selective pressures shaping microbial communities on different evolutionary time scales. It is known that certain traits (*e.g.* salinity preference, methanogenesis) are relatively slow-evolving and thus restricted to only certain lineages, whereas other traits (*e.g.* phage resistance, organic phosphate utilization) can be acquired by a single point mutation or gene acquisition, thus evolving rapidly in response to ecological selection and competition (Martiny *et al.*, 2015). Therefore, habitat filtering might be stronger for slow-evolving traits, while niche partitioning will be more likely for fast-evolving traits.

In this study, we use Hutchinson’s definition of a fundamental niche as the set of abiotic conditions under which an organism can survive and reproduce, and a realized niche as the conditions under which it is actually observed in nature, accounting for both abiotic and biotic (*e.g.* competition, predation, cooperation) interactions (Hutchinson 1957). If two species have identical ecological niches, one should competitively exclude the other (Gause 1934; Tilman 1982) unless competition is weak due to abundant resources. In practice, closely related taxa often compete for space and resources (Cavender-Bares *et al.*, 2009), favouring specialization to reduce overlap in niche space. For example, coexisting (sympatric) taxa within the same genus tend to have different realized niches, experiencing different seasonal growth patterns or responding differently to environmental parameters (Gray *et al.*, 2004; Jaspers and Overmann, 2004; Hunt *et al.*, 2008; Simek *et al.*, 2010; Jezberra *et al.*, 2011; Neuenschwander *et al.*, 2015). Pairs of taxa with similar realized niches can be identified when they co-occur in repeated sampling over space and time, and such co-occurrence networks are readily inferred from deep amplicon sequencing datasets (Friedman and Alm, 2012). Typically, niches are considered as features of species. However, when niches are considered as collections of traits or environmental associations, the Hutchinsonian niche concept can be extended to taxonomic groupings more inclusive than species, even if the biological “reality” of such groups is doubtful. Here, we apply the niche concept to both fine-grained (i.e. sub-genus level) and coarse-grained (*i.e.* genus level) taxonomic units. We focus mainly on niche specialization within genera, showing that specialization is extensive and that lumping bacterial diversity at the genus level obscures finer-scale niche preferences.

Cyanobacteria are widely and naturally present in freshwater ecosystems, and some lineages form blooms under appropriate conditions of temperature and nutrients (Konopka and Broke, 1978; Harke *et al.*, 2016). Several studies have shown that different cyanobacterial genera can co-occur during blooms, thus sharing at least some dimensions of their realized niches (Pearl *et al.*, 2001; Yamamoto, 2009). Nitrogen (N) and phosphorus (P) can both be limiting for bloom formation, and different cyanobacterial taxa apparently have different preferences for N and P concentrations (Dolman *et al.*, 2012). For example, cyanobacteria capable of N-fixation (such as *Dolichospermum*) are associated with more efficient P-utilization at the community level, suggesting P-limitation when N-fixers are abundant (Andersson *et al.*, 2015; Olli *et al.*, 2015). Clearly, N and P utilization are ecologically important traits for cyanobacteria, and may be important for niche partitioning among closely-related strains. In the marine cyanobacterium *Prochloroccocus*, P uptake and metabolism appears to be relatively fast-evolving perhaps due to horizontal gene transfer of P-related genes (Coleman *et al.*, 2010) while light preferences are slow-evolving, and temperature preferences are intermediate (Martiny *et* al., 2015).

We investigated niche partitioning within and between *Microcystis* (*M*) and *Dolichospermum* (*D*), two genera of potentially toxigenic cyanobacteria that bloom nearly every year in a large eutrophic North American lake, Lake Champlain. In a previously described 8-year time-course analysis (2006 to 2013) spanning multiple bloom events, we used 16S amplicon sequencing to broadly survey changes in the lake microbial community over time, generally at the genus level (Tromas *et al.*, 2017). Here, we use Minimum Entropy Decomposition of amplicon sequences (Eren *et al.*, 2014) to identify sub-genus strains (MED nodes; here used interchangeably with “strains”) within each of the two dominant cyanobacterial genera, *M* and *D*, at single nucleotide resolution (*i.e.* each MED node is an exact sequence variant, distinguishable from other variants that differ by at least one nucleotide substitution). MED discards low-entropy nucleotide positions, which effectively filters out sequencing errors at the expense of possibly also removing true polymorphism at very low frequency. Therefore, although MED outputs exact sequences that are actually present in the sample (after denoising), it is possible the MED nodes contain finer-scale genetic variation which could be captured using additional marker genes or whole genome sequencing. A previous study demonstrated that oligotypes (similar to MED nodes) lacked the resolution to distinguish toxic and non-toxic *Microcystis* lineages, but could potentially be informative about other, more phylogenetically conserved niches such as eutrophic (nutrient-rich) vs. oligotrophic (nutrient-poor) lake preferences (Berry *et al.*, 2017). Toxin production is thought to be fast-evolving because toxin biosynthesis genes are widely distributed across cyanobacterial genera, suggesting rapid gain and loss. While horizontal gene transfer is a likely explanation, transfer is probably more frequent among closely-related lineages because toxin gene trees are congruent with ribosomal phylogenies of distantly-related cyanobacteria genera (Rantala *et al.*, 2004).

We used a combination of genetic data (diversity of MED nodes) and matched environmental data (*e.g.* temperature, nutrient concentrations, precipitation) to address three specific questions. First, how similar are the niches of the two dominant cyanobacterial genera *M* and *D*? Second, how similar are the niches of strains within each genus? Third, how does niche similarity change with genetic relatedness? We confirm that *M* and *D* are broadly co-occuring during blooms, but have distinct nutrient preferences. We also identified niche partitioning at the sub-genus level, and observe a general decline in realized niche similarity with genetic distance, consistent with habitat filtering. However, certain niche dimensions (particulate nutrient concentrations) show a complex polynomial relationship with genetic distance, suggesting that a combination of habitat filtering and competitive interactions shapes the evolution of these traits.

## MATERIALS AND METHODS

### Sampling, DNA extraction, purification and sequencing

Open-water season samples (April to November) collected over 8 years (2006 to 2013) from the photic zone of Missisquoi Bay at two different sites (littoral and pelagic) of Lake Champlain, Quebec, Canada (45°02′45′N, 73°07′58″W) were filtered and extracted for DNA sequencing as described in Tromas *et al*., (2017).

### Sequence analysis

A total of 7,949,602 sequences of the 16S rRNA gene V4 region were obtained from 150 lake samples, with a median of 41,982 per sample as previously described (Tromas *et al.*, 2017). These sequences were processed with the default parameters of the SmileTrain pipeline (https://github.com/almlab/SmileTrain/wiki) that includes reads quality filtering, chimera filtering, and merging using USEARCH (version 7.0.1090, http://www.drive5.com/usearch/, default parameter) (Edgar, 2010), Mothur (version 1.33.3) (Schloss *et al.*, 2009), Biopython (version 2.7) and custom scripts. Minimum Entropy Decomposition (MED) was then applied to the filtered and merged reads to partition sequence reads into MED nodes (Eren *et al.*, 2014). MED was performed using the following parameters −M noise filter set to 500 resulting in ∼7% of reads filtered and 941 MED nodes (Data Sheet 2). Samples with less than 1000 reads were removed, yielding a final dataset of 135 samples. Finally, taxonomy was assigned using the assign_taxonomy.py QIIME script (default parameters), and a combination of GreenGenes and a freshwater-specific database (Freshwater database 2016 August 18 release; Newton *et al.*, 2011), using TaxAss (https://github.com/McMahonLab/TaxAss, installation date: September 13^th^ 2016; Rohwer *et* al., 2017). After assignment, nodes that belong to Eukaryotes but still present in the database (Cryptophyta, Streptophyta, Chlorophyta and Stramenopiles orders) were removed, leading to a total of 891 nodes.

### Diversity analysis

Comparing changes in the diversity of *M* strains to the diversity of *D* strains was performed using betta (Willis *et al*., 2017). betta accounts for strains that are present in the environment but not observed in the samples due to incomplete sampling. The number of unobserved strains is estimated based on the number of strains that are observed and their abundances. The total strain diversity was estimated using breakaway (R package v4)which accounts for ecological interactions between strains (Willis and Bunge, 2015).

### Conditionally rare taxa analysis

We investigated the temporal dynamics of *M* and *D* nodes by measuring the composition for each genus in conditionally rare taxa. The matrix of node absolute abundances was used as input for the R script CRT_Functions_v1.1.R (Shade *et al.*, 2014; https://github.com/ShadeLab/ConditionallyRareTaxa) using the default parameters. Conditionally rare taxa are defined as usually-rare taxa that occasionally become very abundant, without showing rhythmic or seasonal patterns.

### Node–environment relationships analysis

To investigate node–environment relationships, we used an environmental data matrix that included: particulate phosphorus in μg/ L (PP, the difference between TP and DP), particulate nitrogen in mg/ L (PN, the difference between TN and DN), total dissolved phosphorus in μg/ L (DP), total dissolved nitrogen in mg/ L (DN), 1-week-cumulative precipitation in mm and 1-week-average air temperature in Celsius. Total nutrients were measured directly from collected lake water and the dissolved nutrients were measured in filtered water (Glass microfiber Filters grade GF/F, 0.7 microns). The detailed measurements of each environment variable are described in Tromas *et al* (2017). In this previous study, we showed that these environmental variables, over the 8 years, were not correlated with one another.

### Response to abiotic factors

To analyze the response of each node to abiotic environmental data, we used a Latent Variable Model (LVM) framework, which combines Generalized Linear Models with Bayesian Markov Chain Monte Carlo (MCMC) methods (boral package in R; Hui 2015, Warton *et al.* 2015). LVM is a model-based approach for analyzing multivariate data (*e.g.* numerous taxa within the response matrix) that partitions the different drivers of taxa co-occurrence patterns into two components: the first is a regression component, which models the taxon-specific environmental responses, and the second is a latent variable component, which is used to identify residual patterns of co-occurrence resulting from unmeasured factors and/or biotic interactions (Letten *et al.* 2015). In this study, we used LVMs to examine how taxa co-respond to abiotic gradients, and used these co-responses as a proxy for niche similarity. To do so, we extracted the *environmental correlation matrix* from the regression model, which, for any two taxa, corresponds to the correlation between their fitted values (x_i_β_j_). In particular, for each node we calculated the predicted probability of mean abundance for each site on linear and nonlinear (*i.e.* Y ∼ X + X^2^) response scales, providing a vector of fitted values. Correlations among the fitted responses of any two taxa were then calculated based upon these vectors (correlating the vector of taxon A with the vector of taxon B, for instance).

We first tested LVMs using a response matrix of two columns, corresponding to the relative abundances of the *D* and *M* genera, relative to the rest of the bacterial community. For these models, *D* and *M* abundances were normalized by the total counts of all bacteria using the centered log ratio to correct for data compositionality (clr-inter genus) (Aitchison, 1986; Paliy and Shankar, 2016) using a a zero-replacement procedure as suggested in Gloor and Reid (2016). To then examine dynamics within these genera, we ran separate LVMs for *D* and *M* nodes (*i.e.* response matrix consisted of the *D* and *M* nodes, respectively). For this second set of LVMs, node abundances were normalized by the total counts of *D* and *M*, respectively, to obtain an intra-genus relative abundance (clr-intra genus) for each node. Environmental variables were standardized to mean zero and unit variance to reduce the correlation between the linear (X) and nonlinear (X^2^) fit, which also simplified model interpretation and stabilized MCMC sampling. As most of the *D* nodes were conditionally rare, we included only the *D* nodes that were present in at least 70% of samples, to avoid overfitting based on too little data. To complement this multivariate LVM analysis and help visualize the univariate response of each genus and node to the suite of environmental variables, we also tested linear (degree-1 polynomial) and nonlinear (degree-2 polynomial) relationships between each response and explanatory variable. We used AIC to find the single best-fit model (either linear or nonlinear) for each relationship and plotted the best-fit model.

Lastly, to test whether co-responses or niche separations were stronger for more closely or distantly related nodes, we extracted the correlations between fitted responses of any two nodes from the *environmental correlation matrix* of each best-fit LVM and examined the relationship between each significant correlation and the pairwise genetic distances of the two respective taxa (percent identity in the V4 region of the 16S gene). Here, significant positive correlations represent co-response between any two taxa whereas significant negative correlations represent the degree of niche separation. The genetic distance between nodes was measured within each genus *M* and *D* using the software MEGA (version 7.0.18) using the p-distance (the proportion of nucleotide sites at which two sequences differ), calculated by dividing the number of sites with nucleotide differences by the total number of sites compared (excluding sites with gaps). R code to reproduce these analyses and relevant figures is provided in Data Sheet 3.

### Co-occurrence analysis

Co-occurrences between *Microcystis* or *Dolichospermum* and other taxa were calculated with SparCC (Friedman and Alm, 2012), with 20 iterations to estimate the median correlation of each pair of MED nodes, and 500 bootstraps to assess the statistical significance. Correlations were then filtered for statistical significance (P < 0.01) and correlations with R > ±0.6, were selected to build networks using Cytoscape (version 3.1.0).

## RESULTS

### Do *Microcystis* and *Dolichospermum* co-occur temporally?

We have previously shown that *M* and *D* were the two most dominant bloom forming cyanobacteria in Lake Champlain’s Missisquoi Bay (Tromas *et al.*, 2017). Both genera were present every year at both littoral and pelagic sampling sites, between 2006 and 2013, with a relative abundance of at least 15% on average during summer (max=24.8%), with the exception of summer 2007 (0.7%) when no substantial blooms occurred (Figure 1). *D* was generally the most dominant genus except for 2009 and 2010 when *M* dominated (Figure S1). *D* and *M* were both present every year, (Figure S1) suggesting that they share similar physico-temporal niche during a bloom, although other aspects of their niches likely differ. For example, *D* can fix nitrogen but *M* cannot (Hajdu *et al*, 2007).

**Figure 1.**
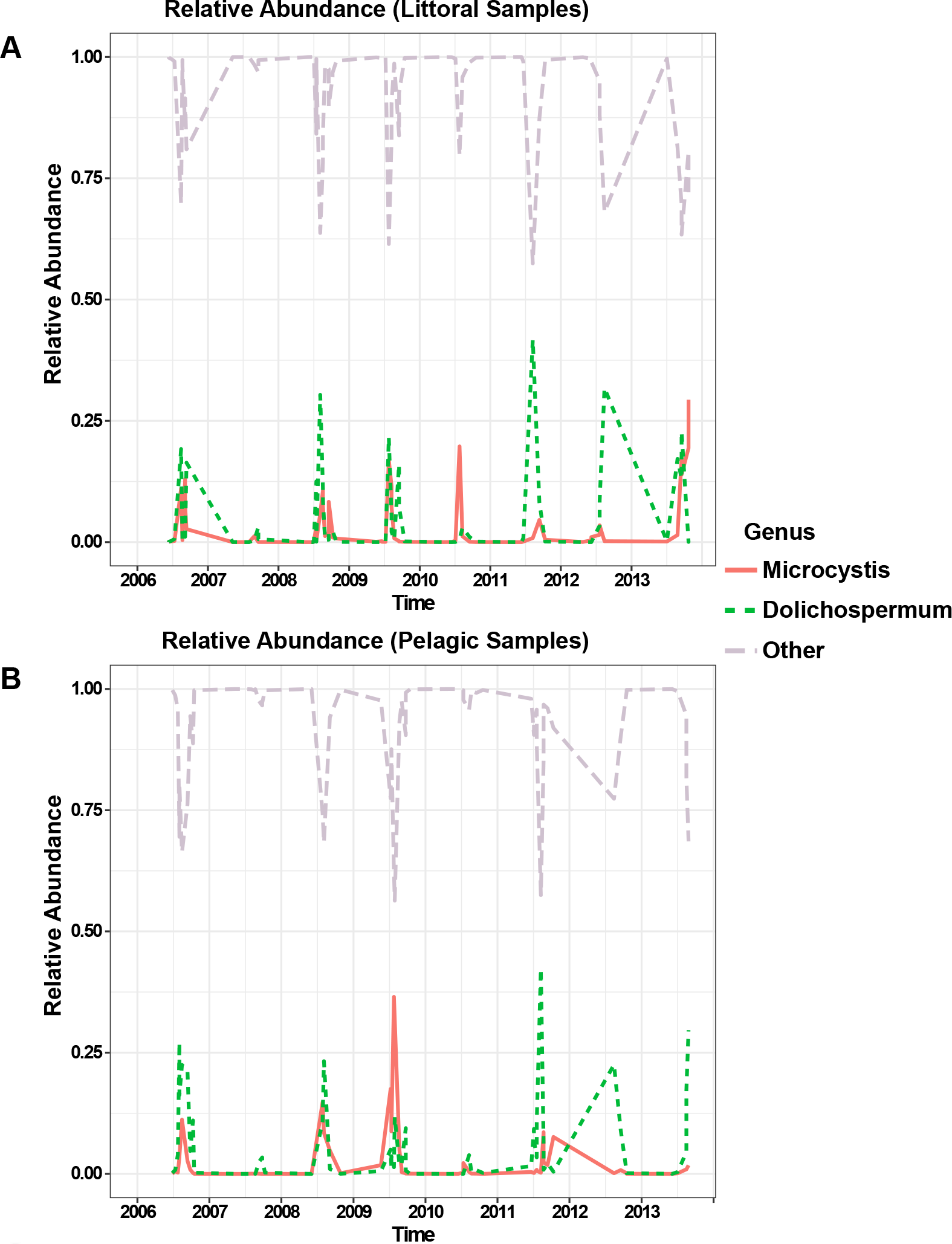
Temporal dynamics of the two dominant cyanobacterial genera over an eight-year time course in littoral. (A) and pelagic (B) sampling sites. Relative abundance of *Microcystis* is shown in solid red, *Dolichospermum* in dashed green and the other members of the bacterial community in dashed light violet. The time scale (x-axis) is in units of years.

### Are *Microcystis* and *Dolichospermum* equally diverse and dynamic?

To explore the temporal dynamics of the finer-scale taxa within each genus, we partitioned each genus into MED nodes, which we call “strains” (Methods). We obtained 25 *D* strains and 6 *M* strains, and after accounting for low-abundance strains that may be missing from the samples using Breakaway (Methods), we conclude that diversity is significantly lower within *M* than within *D* (P<0.001). The difference in the number of nodes is unlikely to be an artefact of sequencing depth, because there is no correlation between the number of sequence reads and the number of nodes per genus (Figure S2). *D* thus appears to be more diverse than *M*.

We then examined whether nodes within the *M* and *D* genera followed the same temporal dynamics. We observed that 15/25 *D* nodes were conditionally rare, meaning that they are usually rare but occasionally become relatively abundant, without following any apparent rhythm (Table 1).

**Table 1.** Conditionally rarity analysis. We compared the proportion of conditionally rare M or D MED nodes to the proportion expected among MED nodes in the entire lake bacterial community using Fisher’s exact test.

|  | Number of conditionally rare nodes | Number of total nodes | Proportion conditional rare / total nodes | Fisher's Exact Test P-value |
| --- | --- | --- | --- | --- |
| **Whole bacterial community** | 95 | 891 | 0.10 |  |
| ***Dolichospermum*** | 15 | 25 | 0.60 | $P < 0.001$ |
| ***Microcystis*** | 1 | 6 | 0.16 | $P > 0.1$ |

The proportion of rare *D* nodes was significantly higher than what is observed in other nodes in the lake community (Fisher′s exact test, *P* < 0.001). Several conditionally rare *D* nodes seemed to dominate several times without showing seasonal patterns (Figure S3). Furthermore, we noticed a shift in node composition after 2011, where node D5505 decreased while D2282 increased in relative abundance. In contrast, we only found one conditionally rare *M* node. However, only two *M* nodes (M5732 and M5733) were consistently dominant over time (Figure S4). Similarly, only two *D* nodes, but not always the same two, dominated at any given time (Figure S3). Overall, these results suggest that *D* is more genetically diverse, and that this genetic diversity varies over time. In contrast, *M* is less diverse and more stable over time.

### Do *Microcystis* and *Dolichospermum* share the same realized niche within the community?

To investigate the niche separation between *D* and *M*, we analyzed their relationships with several environmental conditions measured at the time of sampling (temperature, nutrient concentrations, precipitation and toxin concentrations). We observed that these two cyanobacterial genera have different responses to nutrients (Figure S5) as previously observed (Anderson *et al* 2015; Harke *et al*, 2016). *M* relative abundance was positively correlated with DP (Figure S5; R^2^=7%), in agreement with previous observations (Homma *et al.*, 2008). In contrast, *D* was not significantly correlated with DP, and instead was positively correlated with PP and PN (Figure S5). *D* also responded negatively to dissolved nitrogen, in agreement with previous studies demonstrating that Nostocales (the Order containing *Dolichospermum*) are favoured under conditions of low dissolved inorganic nitrogen, due to their ability to utilize atmospheric nitrogen for growth (Suikkanen *et al*, 2013; Andersson *et al*, 2015). Overall, these results confirmed that *M* and *D* share a spatio-temporal niche during a bloom, but have distinct nutrient preferences.

### How similar are the niches of strains within a genus?

Numerous studies have focused on environmental conditions that favour cyanobacterial blooms, but few studies have examined how different cyanobacterial strains might respond differently to environmental conditions due to niche partitioning. We observed that *D* and *M* strains, within each genus, appear to have qualitatively different dynamics (Figure S3, S4). To test whether these different dynamics were due to niche partitioning, we first analyzed how the different strains within each genus were related to environmental variables, and then used Latent Variable Models (LVMs) to determine the co-responses (niche similarity) as well as niche separation between strains.

We found that nearly all (19/20) of the significant relationships between strain relative abundances (within each genus) and environmental conditions were linear (Figure 2; Table S2). *D* nodes D2282 (red) and D5630 (green) showed opposite responses to nutrients (PP and PN) and precipitation (Figure 2A). The LVMs confirmed a significant niche separation for these niche dimensions (Figure S6). In contrast, nodes D2424 and D5630 both displayed a similar negative relationship (Figure 2A), resulting in a significant co-response (niche similarity) to DN (Figure S6). Within the *M* genus, we observed several nodes with similar responses to PP, PN and DP (Figure 2B), and significant co-responses between nodes were detected for these niches (Figure S7). We also found significant niche separations for DN (involving M5732, M5733 and M5738) and temperature (involving M5734 and M5738) (Figure S7). Overall, the LVM analysis showed that niche separation occurs among nodes within a genus (Figure S8). In *D*, niche separation occurred mainly in the niche dimensions of particulate nutrients (PP and PN) and precipitation (Figure S6), whereas in *M* niche separation occurred mainly according to temperature and DN preferences (Figure S7).

**Figure 2.**
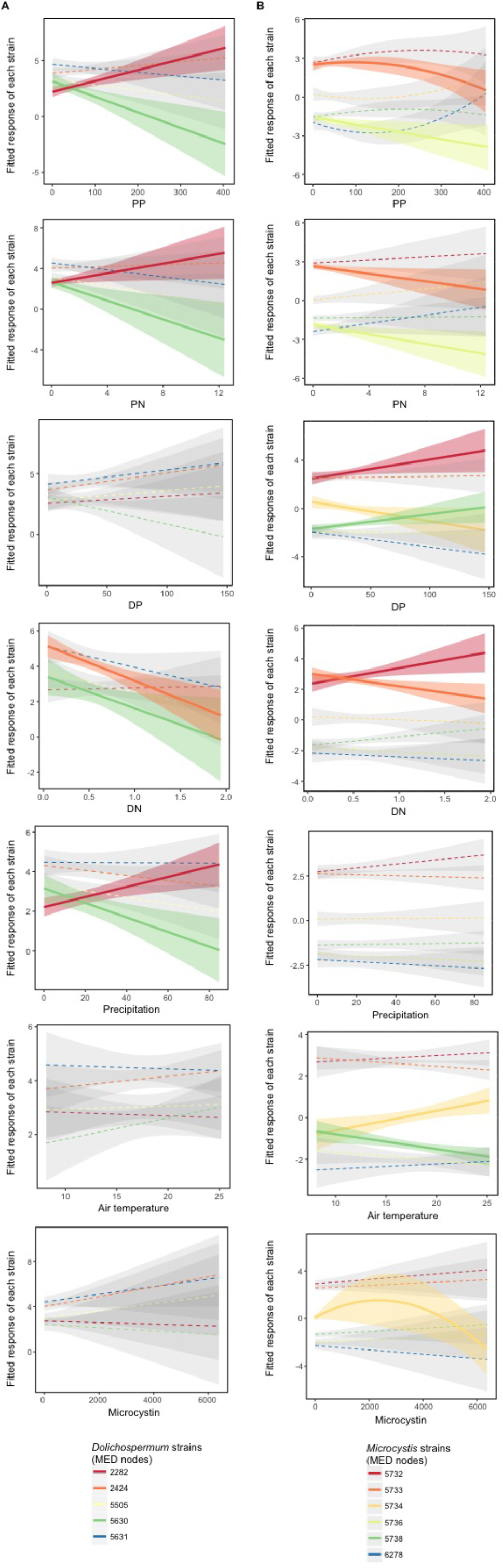
Niche partitioning at the sub-genus level. Best-fit polynomial models of the response of *Dolichospermum* (A) and *Microcystis* (B) nodes to abiotic factors. The relative abundance of each MED node (strain) was computed relative to the total number of reads within each genus using the centered-log ratio (clr) transform. Significant relationships are shown by solid lines and coloured confidence intervals. In most cases, the degree-1 polynomial (linear model) provided the best-fit (see Table S1 for details). For *D*, only the dominant nodes (observed in at least 70% of samples) are shown.

### How do niche preferences vary with genetic distance?

When two taxa share the same realized niche (*i.e.* where they are actually able to survive in the wild, including biotic and abiotic niches), they are more likely to be observed together, and thus to be correlated in survey data. We therefore used co-occurrence patterns between pairs of *D* or *M* nodes as a proxy for similarity in their realized niches, and asked if more genetically similar nodes are more likely to have similar realized niches. Indeed, we found that pairwise SparCC correlation coefficients between nodes tends to decline with genetic distance (Figure 3).

**Figure 3.**
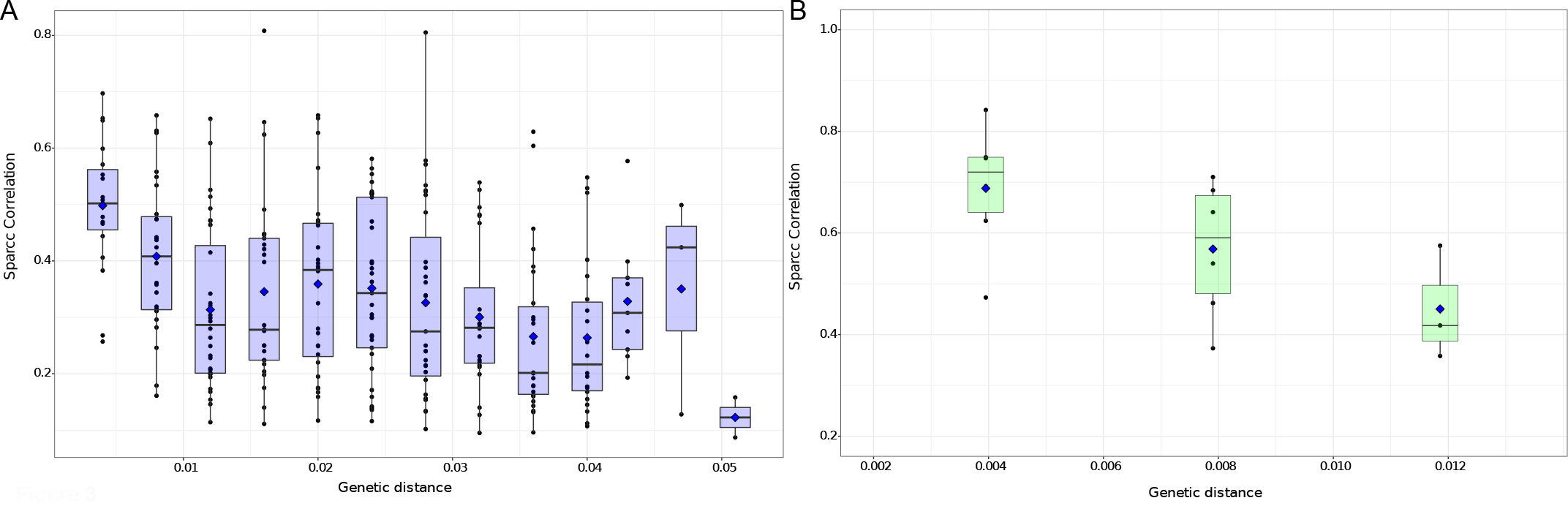
Co-occurrence of strains declines with their pairwise genetic distance. Relationship between co-occurrence (significant SparCC correlation, *P* < 0.05) and genetic distance (p distance) between *Dolichospermum* (A) and *Microcystis* (B) nodes. Blue diamonds represents the mean SparCC correlation for each distance. Boxplots show the median (horizontal line), the 25th and 75th percentile (enclosed in box) and 95% confidence intervals (whiskers). The discreteness observed in the x-axis is due to the discrete number of substitutions in the 16S rRNA gene sequence (*e.g.* exactly 1, 2, 3,… nucleotide differences between pairs).

This pattern was significant within both *D* (linear regression, F(1,279) = 28.3, *P* < 0.001, adjusted R^2^ = 8.9%) and *M* (linear regression, F(1,13) = 7.9, *P* < 0.05, adjusted R^2^ = 33.0%). The higher R^2^ observed for *M* might be explained by its more limited genetic diversity (maximum pairwise genetic distance ∼0.01) compared to *D* (maximum distance ∼0.05). There appears to be a rapid decline in niche similarity as genetic distance goes from 0 to ∼0.01 (adjusted R^2^ = 23.0% in *D* when considering only this distance range) followed by a flatter relationship for genetic distance >0.01.

Ecologically distinct lineages are expected to be associated with distinct surrounding communities, due to a combination of direct microbe-microbe (biotic) interactions and shared preferences for abiotic conditions (Cohan and Koeppel, 2008). We therefore analyzed the co-occurrence patterns between each *D* or *M* node and other bacterial taxa in the lake community. We identified non-cyanobacterial MED nodes that co-occurred with each *D* and *M* node and found that *M* nodes were generally more connected with other members of the bacterial community (Figure S9). We also found that relatively few taxa (4 out of 26; Table S5) are significantly correlated with both *M* and *D*, suggesting that *M* and *D* strains co-occur with distinct sets of other bacteria, and thus have distinct realized niches.

We further investigated whether more closely-related *M* or *D* nodes have more similar correlations with potentially interacting community members. We focused on members of the *Cytophagaceae* family (MED nodes 3667, 5983, and 5984), which are potential predators of cyanobacteria (Rashidan and Bird, 2001), members of the *Rhizobiales* order (nodes 4737, 3705, and 3726), which are potential N-fixers that could provide nitrogen for non-N-fixing *Microcystis* (Louati *et al*, 2015), and all the taxa that co-occurred with both *D* and *M* nodes (nodes 1061, 4674, 7272, and 4756). For each pair of *D* or *M* nodes, we correlated their genetic distance (as in Figure 3) with the absolute difference in the correlations (*r*) with each potentially interacting community member (Figure S11). We found that closely-related *D* nodes tend to have more similar correlations with *Cytophagaceae* node 3667, whereas more distantly-related *D* nodes have more different correlations with node 3667 (Correlation between |Δ*r*| and genetic distance, adjusted-R^2^=0.059, *P <* 0.001). Similar but not statistically significant patterns were observed for several *M* nodes. It is difficult to generalize from these results, but at least in some cases, more phylogenetically similar strains may share conserved interactions with other community members.

Finally, we asked whether the decline in niche similarity with genetic distance (Figure 3) was a common feature of all niche dimensions, or if different abiotic parameters showed different patterns. We considered all significant models, both linear and non-linear as shown in Figure 4 (for *D*) and Figure S10 (for *M).* We found a negative linear relationship between DN niche similarity and genetic distance (adjusted R^2^=25%, *P* = 0.0791). Qualitatively, most other niche dimensions showed a similar pattern for *D* (Figure 4), and among *M* nodes we found a significant negative linear relationship between the correlated fitted response to microcystin concentrations and genetic distance (Figure S10). However, we also identified non-linear relationships between *D* niche similarity and genetic distance for PP (adjusted R^2^=69%, *P* = 0.0184) and PN (adjusted R^2^=63%, *P=* 0.0280). For these niche dimensions, the correlation of fitted responses declines from genetic distances of 0 to ∼0.01, then rises to another peak around ∼0.03 before declining again (Figure 4). While these non-linear models provided significantly better fits to the data compared to linear models, we cannot exclude the possibility that the fits were driven by a few outlying points particular to our data set. Replicating these findings in additional data sets (*e.g.* from different lakes, over different time scales) is therefore essential.

**Figure 4.**
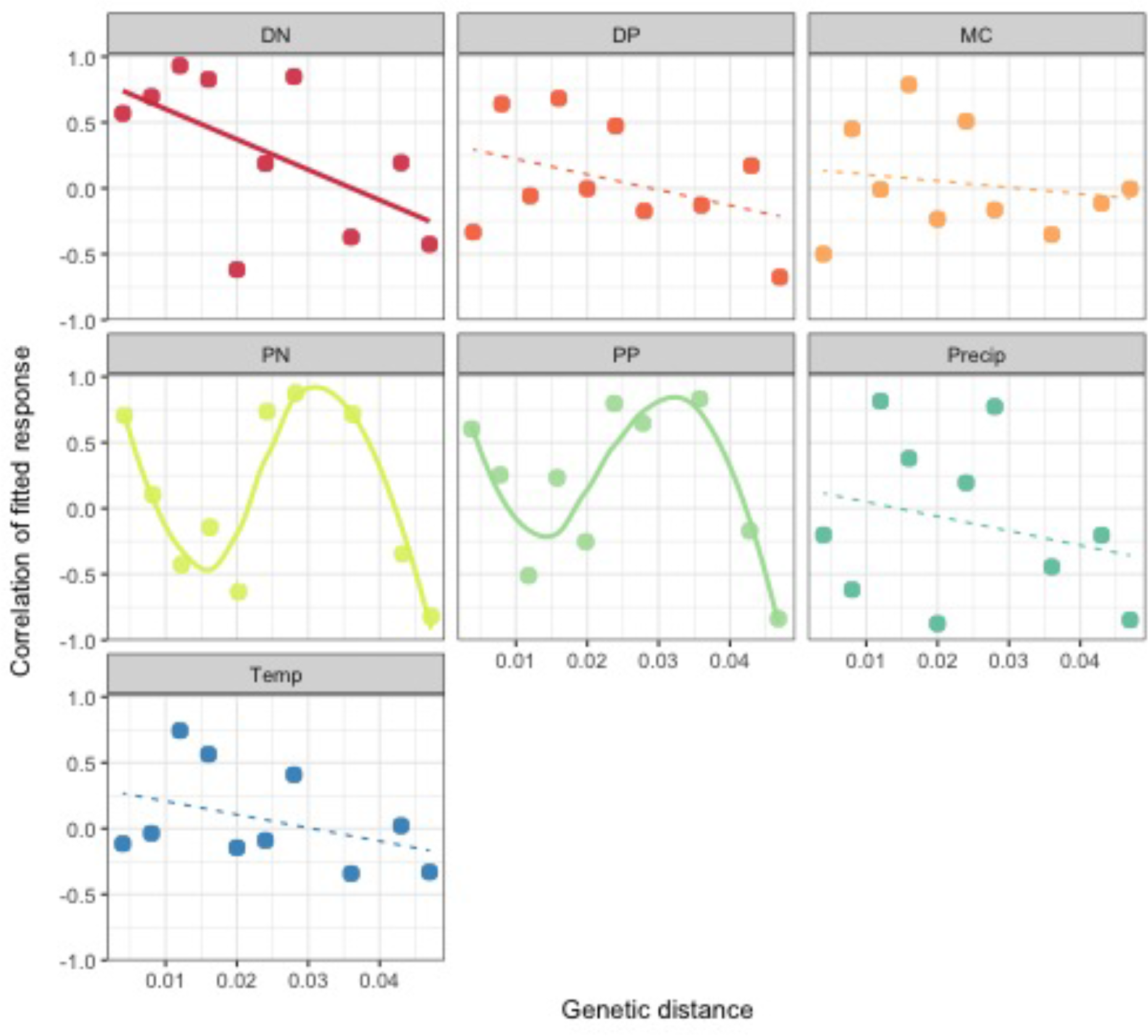
The relationship between genetic distance and the co-response (niche similarity) for *Dolichospermum*. LVMs were used to identify correlations between the responses of MED nodes to each measured environmental parameter (separate panels). Positive correlations of fitted responses indicate similar niches; negative correlations indicate different niche preferences. Genetic distances were computed using the p-distance. Separate model fits were tested with the Akaike information criterion (AIC) for the relationship between each niche dimension and genetic distance. See Table S4 for details of model fits. Significant model fits are shown with thick solid lines; non-significant fits are shown with dashed lines.

## Discussion

Overall, our results show how *Microcystis* and *Dolichospermum* can achieve similar levels of dominance during cyanobacterial blooms by partitioning niche space within and between genera. Over eight years of sampling, we observed that *Dolichospermum* and *Microcystis* generally co-occurred during blooms, suggesting a broadly similar physico-temporal niche (Figure 1; Figure S1). However, we confirmed that *D* was associated with lower concentrations of dissolved nitrogen, consistent with its known ability to fix nitrogen (Harke *et al.*, 2016, Andersson *et al.,* 2015) while *M* was associated with higher concentrations of dissolved phosphorus (Figure S5). These genus-level traits could mask niche differentiation among strains within each genus.

To dissect niche preferences at finer taxonomic resolution, below the genus level, we used minimum entropy decomposition (MED), allowing single-nucleotide resolution of 16S amplicon sequences, *i.e.* each MED node is an exact sequence variant. Finer taxonomic resolution has been shown to increase the power to correctly identify phylogenetic overdispersion, a signature of competitive interactions being more important than habitat filtering (Koeppel and Wu, 2014). However, even at high resolution, the 16S marker gene may be too slow-evolving to be a good marker for fast-evolving traits, such as toxin production (Berry *et al*, 2017) – a trait which likely evolves rapidly by horizontal gene transfer (Moffit and Neilan, 2004).

Despite these limitations, MED analysis provided additional insights into the distinct niches of *M* and *D. Dolichospermum* diversity was higher and most of the MED nodes (strains) were conditionally rare, some of them being bloom-associated in some years but not in others (e.g D2282, Figure S3). *Microcystis* nodes, on the other hand, were less diverse but more consistently bloom-associated (nodes M5732 and M5733; Figure S4) and more correlated with other taxa (Figure S9), suggesting that the two dominant bloom-forming genera might use different ecological strategies, beyond what is already known about nitrogen and phosphorus utilization.

Examining niche partitioning below the genus level also allowed us to detect patterns not evident at the genus level. For example, *Dolichospermum* is associated with higher concentrations of PP and PN at the genus level (Figure S5), but contains a strain (D5630) that shows the opposite pattern (Figure 2, Figure S6). Within *Microcystis*, niche partitioning occurred mainly for temperature and DN, and the two most dominant *M* nodes had a significant niche separation for DN (Figure 2; Figure S7). It is known that *Microcystis* does not fix atmospheric nitrogen, but is able to use refractory N-containing compounds such as urea or amino acids (Moisander *et al*, 2009; Dai *et al*, 2009), and DN is likely needed for toxin production (Monchamp *et al*, 2014). Therefore, different *M* strains could specialize in their preference for different forms of nitrogen.

The relative importance of habitat filtering and competition in shaping microbial communities is widely debated, and distinguishing between the two processes can be technically challenging (Koeppel and Wu, 2014; Cadotte and Tucker 2017). Consistent with a general effect of habitat filtering in selecting for genetically similar cyanobacteria under similar conditions, we found a negative relationship between MED node co-occurrence and pairwise genetic distance (Figure 3). This result is in agreement with an early study showing the importance of habitat filtering in microbial communities (Horner-Devine and Bohannan, 2006) and is also consistent with what was observed by Silverman *et al* (2017) using human microbiome data. Here we considered co-occurrence as a proxy for shared realized niches, including both biotic and abiotic factors.

Consistent with the general importance of habitat filtering in shaping cyanobacterial communities, we observed that closely-related strains tended to have similar co-responses to several measured abiotic environmental parameters. For example, we found negative linear relationships between niche similarity and *Dolichospermum* genetic distance for most niche dimensions, particularly DN (Figure 4). This observation is in agreement with a previous study that showed that bacterial and fungal responses to N fertilization tend to be more similar among close relatives, with a decline in similarity between genetic distances of 0 and 0.05 (Martiny *et al.*, 2015; Amend *et al.*, 2016). In *Microcystis*, we found that more genetically similar strains tended to be observed at more similar concentrations of the toxin microcystin (Figure S10). Although 16S is a poor marker for microcystin production (Berry *et* al., 2017), our results suggest that phylogenetic relatedness is nevertheless somewhat predictive of microcystin concentrations. This means that microcystin production or tolerance is more likely to be shared by close relatives.

In contrast to the general pattern of declining niche similarity with genetic distance, PN and PP both had non-linear relationships with genetic distance in *Dolichospermum* (Figure 4). After an initial decline in co-responses to PN and PP to a genetic distance of ∼0.01, co-response rose to another peak at ∼0.03 before declining again. This response is consistent with near-identical MED nodes sharing the same preferences for PN and PP concentrations, and that competition between close-relatives (up to genetic distance of ∼0.01) imposes divergent selection for distinct niche preferences. The similarity in PN and PP niches for more distant relatives (distance ∼0.03) can be explained if distant relatives have diverged to reduce competition in other niche dimensions, allowing them to converge in PN and PP preferences. It is unclear why this non-linear relationship is observed for PN and PP, but not for other niche dimensions. One possibility is that P uptake and metabolism genes are easily acquired by horizontal gene transfer, as observed in marine cyanobacteria (Coleman *et* al., 2010) and may thus be more rapidly evolving, resulting in non-linear relationship with phylogenetic distance. Yet it is unclear why the non-linear pattern is observed for particulate but not dissolved N and P. Particulate nutrients could be a marker for biomass (e.g. bloom density), but it is equally unclear how rapidly “bloom preference” would be expected to evolve. The same non-linear pattern might be expected for microcystin, due to frequent gain/loss of the underlying biosynthetic genes. However, we observe a linear relationship (Figure S10) between microcystin concentrations and genetic distance among *Microcystis* strains – but only in a very narrow range of genetic distance (99-100% identity). Therefore, non-linear relationships could become apparent among more distantly related strains (e.g. 95-99% identity).

To investigate how biotic niches change over evolutionary time, we studied how cyanobacterial interactions (SparCC correlations) with non-cyanobacterial community members varied with genetic distance among cyanobacteria. We found that genetically similar *Dolichospermum* strains tended to have similar interactions with a strain of *Cytophagaceae*, a potential cyanobacterial predator, whereas more distantly related *Dolichospermum* strains diverged in terms of their interactions (Figure S11). We limited our analyses to only the most common interacting partners (common to *M* and *D*) or those previously suspected to interact with cyanobacteria via predation or cross-feeding. Future studies could expand on these analyses in a more comprehensive fashion to quantify how biotic interactions evolve over different time scales and in different lineages.

Overall, our results show how different traits may have different relationships with genetic distance, highlighting the importance of considering each niche dimension separately, because adaptation to different niche dimensions can occur at dramatically different rates (Martiny *et al.*, 2015). Our results also suggest that the same trait could evolve at different rates in different cyanobacterial lineages. For example, temperature adaptation evolves at an intermediate rate in *Prochloroccocus* (Martiny *et al.*, 2015) and perhaps at a slower rate in hotspring *Synechococcus* (Becraft *et al.*, 2011; Becraft *et al.*, 2015). Using our comparative framework of two broadly sympatric genera, we identify a general decline in ecological similarity with genetic distance – although certain traits (*e.g.* PN and PP in *Dolichospermum* but not in *Microcystis*) go against this trend. A future challenge in microbial ecology and evolution will be to determine which traits are generally fast- or slow-evolving, and most interestingly, which lineages provide exceptions. For example, particularly rapid evolution of a generally slow-evolving trait in a particular lineage could provide evidence for an unusual genetic architecture or strong selective pressure on that particular trait in that particular lineage.

## Data availability

Raw sequence data have been deposited NCBI GenBank under BioProject number PRJNA353865: SRP099259.

## Conflict of Interest

The authors declare no conflict of interest.

## Acknowledgments

We thank Francis Hui, Andrew Letten and David Warton for advice on the LVM analyses, and the two peer reviewers for their constructive comments that improved the manuscript. We also thank everyone who participated in sampling, data collection and analysis. This research was funded by a Natural Sciences and Engineering Research Council (NSERC) Discovery grant and a Fonds de Recherche du Québec Nature et Technologies (FRQNT) New Researcher grant to BJS, a FRQNT Programme de recherche en partenariat sur les cyanobactéries grant to CWG, and the federal government interdepartmental Genomics Research and Development Initiative (GRDI). NT is funded by a project from the European Union’s Horizon 2020 research and innovation program under the Marie Sklodowska-Curie grant agreement No 656647.

